# AGO2a but not AGO2b mediates antiviral defense against the infection of wildtype Cucumber mosaic virus in tomato

**DOI:** 10.1101/2022.11.23.517585

**Authors:** Li-Ling Zhao, Ying-Fang Chen, Xing-Ming Xiao, Hai-Ying Gao, Jia-Min Cao, Zhong-Kai Zhang, Zhongxin Guo

## Abstract

Evolutionarily conserved antiviral RNA interference (RNAi) mediates a primary antiviral innate immunity preventing the infection of broad spectrum viruses in plants. However, the detailed mechanism in plants is still largely unknown, especially in important agricultural crops including tomato. On the other aspect, varieties of pathogenic viruses evolve to possess Viral Suppressor of RNA silencing (VSR) to suppress antiviral RNAi in host. Due to the prevalence of VSR, it is still skeptical that antiviral RNAi truly functions to prevent the invasion of natural wildtype viruses in plants and animals. In the research, it is for the first time we applied CRISPR-Cas9 to generate *ago2a, ago2b* or *ago2ab* mutants for two differentiated *Solanum lycopersicum* AGO2, one key effector in antiviral RNAi. We found that AGO2a but not AGO2b was significantly induced to inhibit the propagation of not only VSR-deficient Cucumber mosaic virus (CMV) but also wildtype CMV-Fny in tomato, however, both AGO2a and AGO2b did not regulate disease induction after the infection of either virus. Our findings firstly reveal a prominent role of AGO2a in antiviral RNAi innate immunity in tomato and demonstrate that antiviral RNAi evolves to defend the infection of natural wildtype CMV-Fny in tomato, however AGO2a-mediated antiviral RNAi does not play major roles in promoting tolerance of tomato plants to CMV infection for maintaining health.

## INTRODUCTION

RNAi is evolutionarily conserved in eukaryotes, it essentially regulates varieties of biological processes in organisms (Chinnusamy and Zhu, 2009; Bologna and Voinnet, 2014; Iwasaki et al., 2015; Li and Wang, 2019). Antiviral RNAi is a fundamental antiviral innate immunity in plants and animals, it plays vital roles to prevent host from the infection of all kinds of viruses (Ding, 2010; Guo et al., 2019a; Jin et al., 2022; Lopez-Gomollon and Baulcombe, 2022). The core pathway of antiviral RNAi in plants has been proposed with the identification of several key components, mainly based on research in model plant Arabidopsis (Gong et al., 2022; Zhao and Guo, 2022). In Arabidopsis, after viral infection the double-stranded viral RNA replicate intermediate will be perceived and processed into 21-24bp duplex viral small inference RNAs (vsiRNAs) by different Dicer-like proteins (DCLs), which will produce primary vsiRNAs (Garcia-Ruiz et al., 2010). The primary duplex vsiRNAs will be loaded into effector Argonaute proteins (AGOs) to form RNA-induced silencing complex (RISC) (Gu et al., 2012; Carbonell and Carrington, 2015; Zhang et al., 2015; Niaz, 2018). The passage strand of duplex vsiRNA will be cleaved and then mature RISC will target complementary viral RNAs through the guide strand of vsiRNA, then AGOs in RISC will mediate the degradation or inhibit the transcription of translation of viral RNAs through post translational gene silencing (PTGS) or transcriptional gene silencing (TGS) to restrict viral infection (Matzke and Mosher, 2014; Fang and Qi, 2016; Guo et al., 2019a). In the process, adequate secondary vsiRNA has to been produced through RNA-dependent Polymerases (RDRs) by templating the erratic viral RNA to ensure efficient antiviral innate immunity (Wang et al., 2010; Wang et al., 2011; Sanan-Mishra et al., 2021). A new class of virus-activated endogenous siRNA (vasiRNA) dependent of RDR1 was discovered in Arabidopsis, it may confer another layer of antiviral RNAi innate immunity to plant (Cao et al., 2014; Borges and Martienssen, 2015).

Notably, during the arm-race between host and virus, viruses evolve to possess viral suppressor of RNA silencing (VSR) to disturb antiviral RNAi at distinct steps in the pathway and function as the key factor of viral virulence and pathogenesis (Burgyán and Havelda, 2011; Jamous et al., 2011; Derrien et al., 2018). The prevalence of VSRs and their potent inhibition function on antiviral RNAi have seriously hindered our appreciation of the antiviral immunity. Now it is still questioned whether antiviral RNAi functions to counter the infection of wildtype viruses in nature. On the other hand, the existence of VSR also seriously hinders our effort to identify novel components in antiviral RNAi through genetic screen based on wildtype viruses. It can be found that most of known components in the antiviral RNAi, such as DCLs, AGOs and RDRs, were identified based on their shared functions in silencing transgenes or endogenous genes (Baulcombe, 2004; Ding, 2010; Willmann et al., 2011; Carbonell and Carrington, 2015), an applicable genetic screen for identifying specific components in antiviral RNAi is not available until an effective genetic screen to identify *antiviral RNAi-defective* (*avi*) Arabidopsis mutant was developed recently through VSR 2b-deficient Cucumber mosaic virus (CMV-Δ2b) (Guo et al., 2017; Zhu et al., 2017; Gao et al., 2018; Guo et al., 2018). Now the detailed mechanism of antiviral RNAi in plants is still unclear, especially in important agricultural crops.

Tomato is one of the most important agricultural crops in the world, valued at 102.6 billion US dollars in 2020, with yield estimated at 186.8 million tons in 2020 (FAOSTAT, 2022), it is often threatened by varieties of pathogenic plant viruses (Xu et al., 2017; Rivarez et al., 2021). It has been found that some known key components of antiviral RNAi machinery, such as DCLs, AGOs and RDRs are conserved in the genome of tomato. However, their antiviral functions have not been systematically studied. Notably, tomato evolves to possess multiple differentiated homologs for some of these key components including AGO2 which AGO2a and AGO2b, two tandem repeated homologs, are evolved in tomato (Bai et al., 2012). AGO2 is one of key effectors forming RISC with 21nt or 22nt siRNAs to specifically mediate antimicrobial defense in plants (Zhang et al., 2011; Guo et al., 2019a). AGO2 is found to defend against the infection of different species of viruses through antiviral RNAi in plants and animals (Ding and Voinnet, 2007; Guo et al., 2019a). In plants, it has been reported that AGO2 in Arabidopsis (AtAGO2) can limit the infection of CMV, Turnip mosaic virus (TuMV), Potato virus X (PVX), Turnip crinkle virus (TCV) in Arabidopsis (Wang et al., 2011; Carbonell et al., 2012; Garcia-Ruiz et al., 2015; Brosseau and Moffett, 2015; Zheng et al., 2019; Brosseau et al., 2020), and AGO2 homolog in *Nicotiana benthamiana* (NbAGO2) was also found to defend against the infection of wildtype TuMV, PVX, TCV, Tomato ringspot virus (ToRSV), Sweet potato mild mottle virus (SPMMV) and Tomato bushy stunt virus (TBSV) in *N. benthamiana* (Scholthof et al., 2011a; Odokonyero et al., 2015; Paudel et al., 2018; Diao et al., 2019; Kenesi et al., 2021), though AGO2 homolog in rice (OsAGO2) has been reported to increase plant susceptibility to Rice black-streaked dwarf virus (Wang et al., 2021). Intriguingly, it was found that AtAGO2 or NbAGO2 can only prevent Arabidopsis or *N. benthamiana* from the infection of CMV-Δ2b but not wildtype CMV-Fny (Wang et al., 2011; Ludman et al., 2017). However, the antiviral function of both AGO2a and AGO2b in tomato is still elusive and needs to be clarified with true knockout mutants (Kwon et al., 2020).

CMV is an economically important plant pathogenic virus in the family *Bromoviridae* to infect over 1200 plant species including important crops including tomatoes (Palukaitis et al., 1992; Scholthof et al., 2011b). It is also a model plant virus for studying the interaction between host plant and virus (Palukaitis and García-Arenal, 2003; Jacquemond, 2012; Salanki et al., 2018). CMV genome is composed of three single-stranded positive-sense RNAs which encode five viral proteins, helicase 1a protein, RNA dependent RNA Polymerase 2a protein, Movement Protein (MP), Coat Protein (CP) and VSR 2b protein (Palukaitis and García-Arenal, 2003; Mochizuki and Ohki, 2012). In previous research, we found that wildtype CMV-Fny can infect wildtype Arabidopsis plants or antiviral RNAi-defective Arabidopsis mutants and induce similar disease symptoms in these plants with comparable viral accumulation, due to the potent inhibition of 2b on antiviral RNAi. However, CMV-Δ2b can only abundantly accumulate and induce disease symptoms in antiviral RNAi-defective Arabidopsis mutants, but it cannot efficiently infect to cause disease in wildtype Arabidopsis which antiviral RNAi is intact (Wang et al., 2011; Guo et al., 2017; Guo et al., 2018). Based on these findings, we established a robust platform to study antiviral RNAi in Arabidopsis through VSR-deficient CMV (CMV-Δ2b) (Guo et al., 2017; Guo et al., 2018; Zhao and Guo, 2022). Excitingly, we found that tomato is also a natural host of CMV. Therefore, CMV-Δ2b together with wildtype CMV could also provide a powerful tool to dissect antiviral RNAi in tomato.

In the research, we utilized CRISPR to generate *ago2a, ago2b* single or *ago2ab* double knockout mutants in our effort to dissect antiviral RNAi immunity in tomato with CMV-Fny and CMV-Δ2b. It was found that *ago2a* but not *ago2b* displayed increased viral accumulation after the infection of either CMV-Fny or CMV-Δ2b, indicating that AGO2a but not AGO2b prevented the infection of not only CMV-Δ2b but also wildtype CMV in tomato. Surprisingly, *ago2a, ago2b* or *ago2ab* did not show developmental defects or difference in disease symptom compared to wildtype tomato after the infection of either virus, indicating they did not regulate plant development or disease symptom induction in tomato. We further found that AGO2a but not AGO2b was significantly induced after viral infection, and only AGO2a protein can be readily detected after transient expression, which may underlie their distinct function in antiviral immunity. Thus, in the research we developed an effective platform to study antiviral RNAi in tomato through CMV and its mutant variant, and our findings not only firstly reveal a prominent role of AGO2a in antiviral RNAi innate immunity but also demonstrate that antiviral RNAi evolves to defend the infection of natural wildtype CMV in tomato.

## RESULTS

### 1. Generation of *ago2a, ago2b* or *ago2ab* tomato mutants through CRISPR

AGO2a and AGO2b are respectively encoded by two tandem repeated homologs Solyc02g069260 and Solyc02g069270 in tomato. They are respectively composed of 1042 or 977 amino acids in full-length, and both contain five different domains: ArgoN, DUF1785, PAZ, ArgoL2 and Piwi from N-terminal to C-terminal (Fig S1A). AGO2a and AGO2b show high identity (80%) to each other (Fig. S1B and Fig S2), both AGO2a and AGO2b also show high identity (over 70%) to NbAGO2 (Fig. S1B and Fig S2), although a specific 20 aa region in the ArgoN domain is missed in AGO2b compared to AGO2a or NbAGO2 or other AGO2 proteins (Fig S3). However, both AGO2a and AGO2b show high divergence (∼80%) to AtAGO2 or to OsAGO2 (∼100%) (Fig. S1B and Fig S2), and they lack the uncharacterized domain on the N-terminal of AtAGO2 or OsAGO2 (Fig S1A), suggesting that tomato AGO2a and AGO2b may function differently in antiviral RNAi compared to AtAGO2 or its homologs in other plants.

We then utilized CRISPR-Cas9 to generate *ago2a, ago2b and ago2ab* knockout mutants for the first time in order to characterize their function in antiviral RNAi in tomato, since AGO2a or AGO2b knockout mutants are still not available. For generating *ago2a* or *ago2b* single mutant, we designed two specific CRISPR guide RNAs targeting two different protospacer adjacent motif (PAM) sites on the 1^st^ exon of *AGO2a* (TGG^26-28^ and TGG^87-89^) (Fig 1B) or 1^st^ exon *AGO2b* (TGG^207-209^ and TGG^261-263^) (Fig 1E). For generating *ago2ab* double mutant, we designed two specific CRISPR guide RNAs which respectively targeted two different PAM sites on 2^nd^ exon of *AGO2a* (AGG^3390-3409^) or 1^st^ exon of *AGO2b* (TGG^24-26^) (Fig 1H). These CRISPR guide RNAs were respectively constructed into vector pHEE401, in which Cas9 was integrated, to produce three different expression vectors, pHEE401-AGO2a, pHEE401-AGO2b, pHEE401-AGO2ab (Fig 1A). They were further respectively transformed into tomato Micro-Tom using Agrobacterium-mediated method to edit target genes. We successfully obtained multiple transgenic lines for each transformation, then sequenced the targeting regions to identify homozygotic mutants, in which CAS9 was also segregated out, in T2 generation transgenic plants.

**Figure 1.**
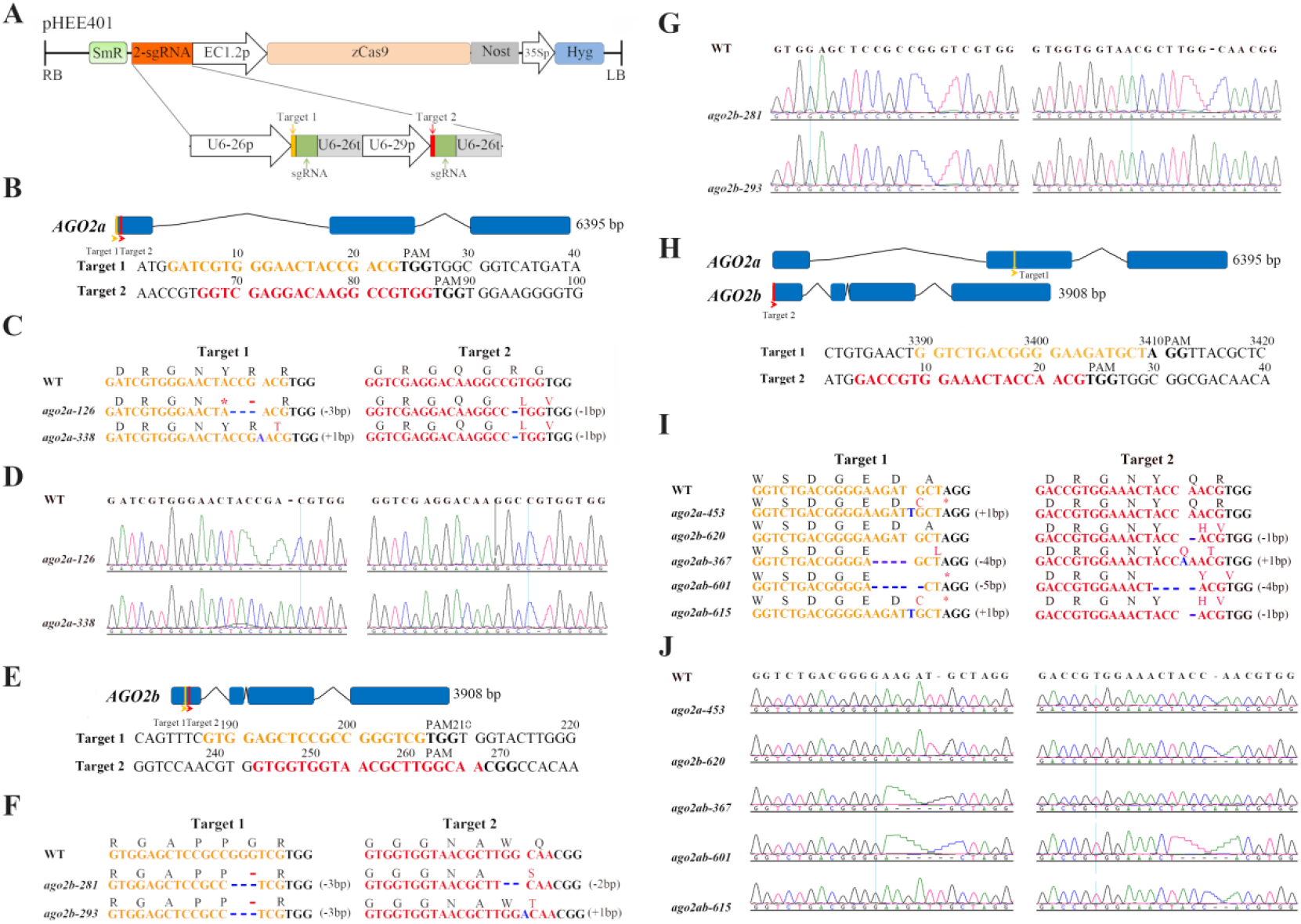
CRISPR/Cas9-mediated *AGO2a, AGO2b* and *AGO2ab* mutations in Micro-Tom. (**A**) Schematic diagram to illustrate the construction of pHEE401 binary vector, used for editing *AGO2a, AGO2b or AGO2ab* gene in Micro-Tom. RB/LB, T-DNA right/left border; U6-26p and U6-29p, two Arabidopsis U6 gene promoter; U6-26t, U6-26 terminator with downstream sequence; EC1.2p, EC1.2 promoter; zCas9, Zea mays codon-optimized Cas9; Nost, *nos* gene terminator; 35Sp, CaMV 35S promoter; Hyg, hygromycin-resistance gene; SmR, Streptomycin-resistance gene. In the structure of sgRNA, target (19 or 20-bp in length) was marked with yellow or red while the green part represents 76-bp sgRNA scaffold. Schematic illustration of two targets in *AGO2a* gene (**B)**, *AGO2b* gene (**E)** and *AGO2a/b* gene (**H)**. Exons are indicated by rectangles and introns indicated by lines. Positions of two small guide RNA (sgRNA) targets are indicated with orange and red arrowheads, respectively. Target sequences are highlighted with the same color as the target. PAM is indicated with bold font. (**C, F and I)** Mutations in *AGO2a and AGO2b* alleles of the *ago2a-126/338*/*453, ago2b-281*/*293/620 and ago2ab-367*/*601*/*615* transgenic lines. Sequences and encoded amino acids of sgRNA target regions are shown. Sequence changes by mutation are marked in blue, while amino acid changes in red. (**D, G and J**) Sanger sequencing results of mutations. DNA fragments around the target sequences were amplified by PCR and then subjected to sequencing analysis.

After confirmation by Sanger-sequencing, we respectively obtained three allelic *ago2a, ago2b* or *ago2ab* mutants. Among them, *ago2a-126* or *ago2a-338* respectively contains 3bp and 1bp deletion, or 1bp insertion and 1bp deletion in target 1 region and target 2 region on *AGO2a* induced by pHEE401-AGO2a (Fig 1C-D), *ago2a-453* only contains 1bp insertion in target 1 region on *AGO2a* induced by pHEE401-AGO2ab (Fig 1I-J). Mutant of *ago2b-281* or *ago2b-293* respectively contains 3bp and 2bp deletion, or 3bp deletion and 1bp insertion in target 1 and target 2 on *AGO2b* induced by pHEE401-AGO2b (Fig 1F-G), *ago2b-620* only contains 1bp deletion in target 2 region on *AGO2b* induced by pHEE401-AGO2ab (Fig 1I-J). However, *ago2ab* double mutants *ago2ab-367, ago2ab-601* or *ago2ab-615* respectively contained 4bp deletion and 1bp insertion, 5bp and 4bp deletion, or 1bp insertion and 1bp deletion in target 1 on *AGO2a* and in target 2 on *AGO2b* induced by pHEE401-AGO2ab (Fig 1I-J). All these small In-Del mutations were localized in exons of AGO2a or AGO2b and caused frameshift mutation on *AGO2a* or *AGO2b* genes. Therefore, *ago2a, ago2b* or *ago2ab* knockout mutants were successfully generated through CRISPR for further research.

### 2. Antiviral immunity is compromised in tomato *ago2a* knockout mutants

To find out AGO2a function in antiviral defense in tomato, we then respectively infected wildtype Micro-Tom and *ago2a-126, ago2a-338* and *ago2a-453* mutants with wildtype CMV-Fny or CMV-Δ2b. We found all mock *ago2a* mutants did not show developmental defects compared to wildtype Micro-Tom (Fig 2A and Fig S4), indicating AGO2a does not regulate plant growth and development. After CMV-Δ2b infection, these *ago2a* mutants did not show visible developmental defects either (Fig 2A). However, about 20 days after the infection of wildtype CMV-Fny, both wildtype Micro-Tom and all three *ago2a* mutants developed severe disease symptoms, such as small statue, mosaic lesions and curly leaves (Fig 2A and S5A). These results indicate that VSR 2b of CMV is critical to induce disease symptoms in tomato after viral infection. However, *ago2a* mutants did not show enhanced disease symptoms in tomato after the infection of CMV-Δ2b or CMV-Fny, indicating AGO2a did not function importantly in preventing disease induction after viral infection in tomato.

**Figure 2.**
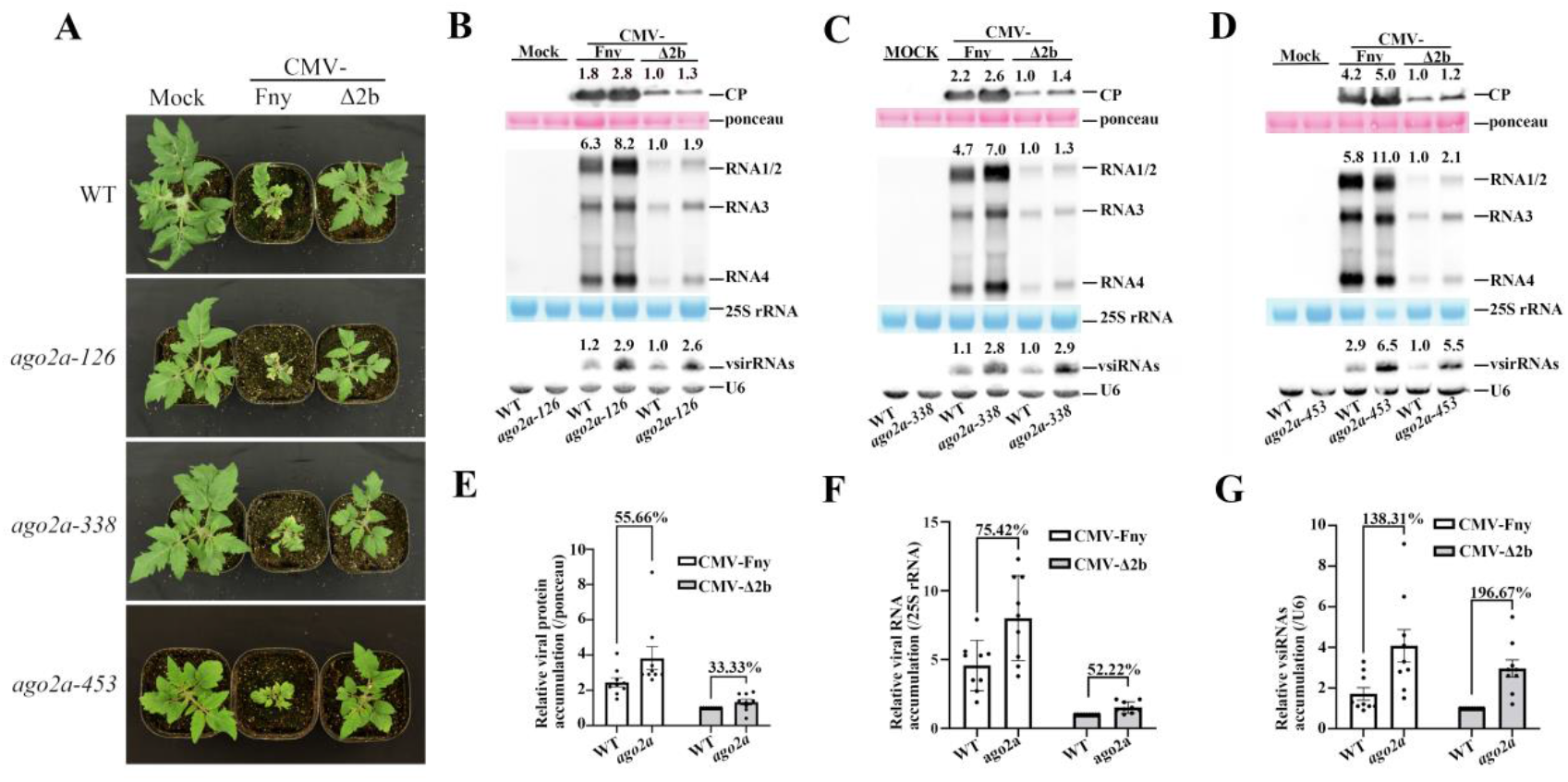
Antiviral immunity is compromised in tomato *ago2a* knockout mutants. (**A**) Wild-type (WT) and three lines of *ago2a* mutants (*126, 338* and *453*) were photographed 19 days post-inoculation (dpi) with buffer (Mock), CMV-Fny or CMV-Δ2b. Symptoms induced with CMV-Fny were shown in Figure S5**A**. (**B**-**D**) Viral protein, viral RNAs and vsiRNAs accumulation in WT and three lines of *ago2a* plants, infected with CMV-Fny or CMV-Δ2b at 19 dpi. Values at the top panel represent corresponding hybridization signal intensity of viral CP, RNAs 1-4, and vsiRNAs. The intensity of WT inoculated with CMV-Δ2b was set as 1. Ponceau, 25S rRNA and U6 RNA were used as the loading controls. Blot-experiments per mutant line were repeated three times with similar results showed in Figure S5 (**B**-**G**). Relative viral protein accumulation (**E**), viral RNAs accumulation (**F**) and vsiRNAs accumulation (**G**) in WT and *ago2a* plants infected with CMV-Fny or CMV-Δ2b at 19 dpi. Values represent the average of nine individual blot-experiments results, three lines of *ago2a* with three technical replicates each. Error bars represent SD. ‘‘%’’ indicates the percentage of increased accumulation in *ago2a* compared to WT (100%).

To further find whether viral accumulation was affected in *ago2a* mutants, we respectively examined viral CP protein or viral genomic RNAs in each of three allelic *ago2a* mutants by Western blot or Northern blot. It was found that CP protein level was clearly increased in all three *ago2a* mutants after either CMV-Fny or CMV-Δ2b infection compared to Wildtype Micro-Tom plants (Fig 2B-D up and S5B-G up). Consistently, viral genomic RNAs accumulation were also dramatically increased in all three allelic *ago2a* mutants compared to Wildtype Micro-Tom plants after the infection of either wildtype CMV-Fny or CMV-Δ2b (Fig 2B-D middle and S5B-G middle). Northern blot analysis showed that the accumulation of viral siRNAs (vsiRNAs) in *ago2a* mutants was also increased compared to wildtype Micro-Tom plants (Fig 2B-D down and S5B-G down), indicating vsiRNA biogenesis was not affected in *ago2a* mutants.

To further demonstrate accumulation variation of CP, viral RNAs and vsiRNAs in *ago2a* mutant, we statistically calculated their relative accumulation based on three replicate results for each different *ago2a* mutant. It was found that, after the infection of CMV-Fny or CMV-Δ2b, viral CP was significantly increased by ∼55% or ∼33% in *ago2a* mutant compared to wildtype Micro-Tom (Fig 2E), viral RNAs was significantly increased by ∼75% or ∼52% (Fig 2F), and vsiRNAs was significantly increased by ∼138% or ∼196% (Fig 2G). These results indicate that AGO2a not only defends against the infection of CMV-Δ2b but also functions to restrict the infection of wildtype CMV virus in tomato. Surprisingly, this result is different from findings in Arabidopsis in which wildtype CMV-Fny is efficiently propagated not only in wildtype Arabidopsis but also in antiviral RNAi-defective Arabidopsis plants because 2b potently inhibits antiviral RNAi in wildtype Arabidopsis. However, AGO2 does not directly regulate vsiRNAs biogenesis in tomato, just like in Arabidopsis.

### 3. AGO2b does not function in antiviral immunity in tomato

To further investigate the antiviral function of AGO2b in tomato, we also infected *ago2b-281, ago2a-293* and *ago2b-620* mutants along with wildtype Micro-Tom with wildtype CMV-Fny or CMV-Δ2b, respectively. We found that all three mock *ago2b* mutants did not exhibit any visible defects in growth and development (Fig S4). And after CMV-Δ2b infection, none of three allelic *ago2b* mutants display observable disease symptoms compared to wildtype Micro-Tom (Fig 3A). However, after the infection of CMV-Fny, these *ago2b* mutants developed typical disease symptoms like wildtype Micro-Tom plants (Fig 3A and Fig S6A). Thus, like AGO2a, AGO2b do not regulate plant growth and development in tomato either, and neither AGO2a or AGO2b function importantly to prevent disease symptom induction in tomato after viral infection.

**Figure 3.**
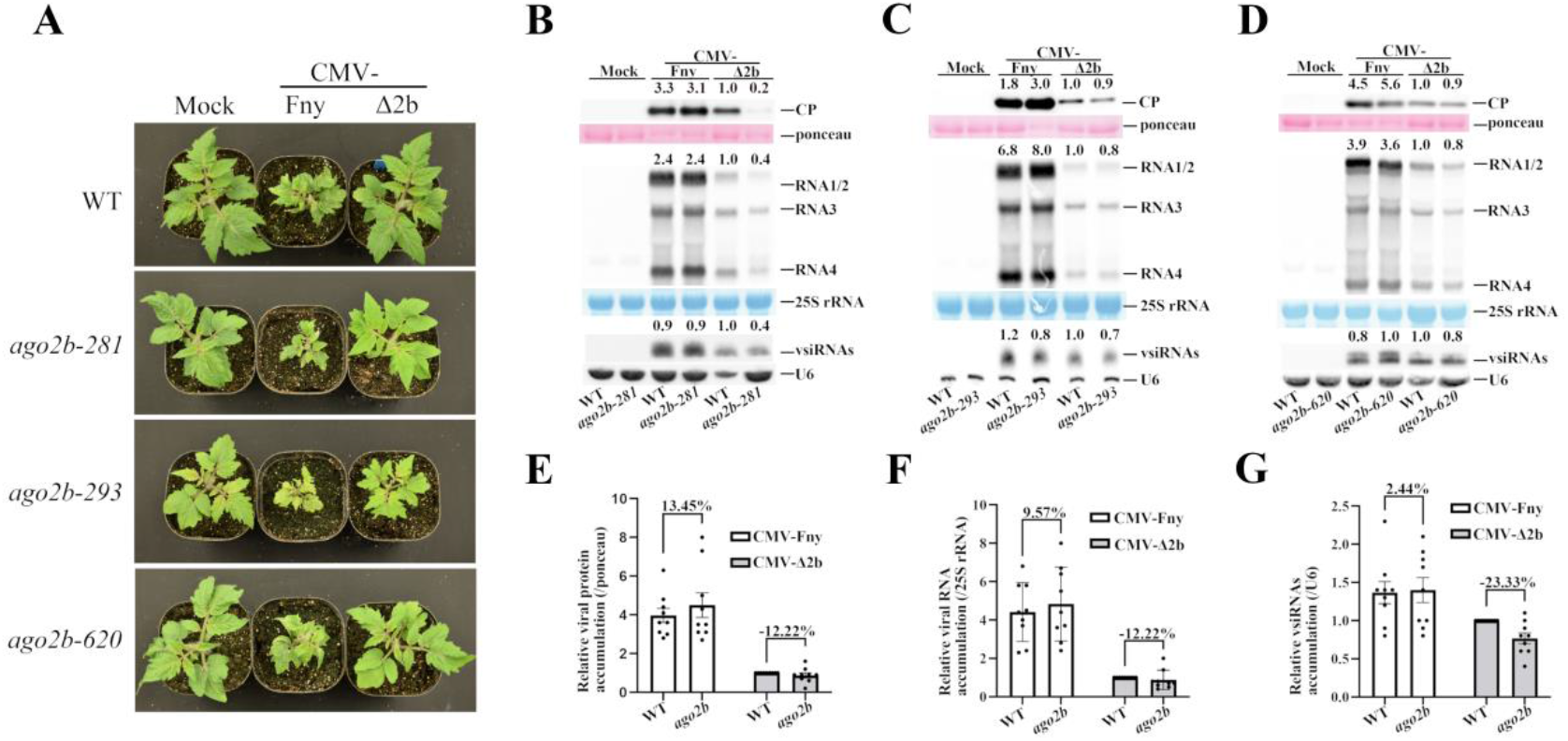
AGO2b does not function in antiviral immunity in tomato. (**A**) Wild-type (WT) and three lines of *ago2b* mutants (*281, 293* and *620*) were photographed 19 days post-inoculation (dpi) with buffer (Mock), CMV-Fny or CMV-Δ2b. Symptoms induced with CMV-Fny were shown in Figure S6**A**. (**B**-**D**) Viral protein, viral RNAs and vsiRNAs accumulation in WT and three lines of *ago2b* plants, infected with CMV-Fny or CMV-Δ2b at 19 dpi. Values at the top panel represent corresponding hybridization signal intensity of viral CP, RNAs 1-4, and vsiRNAs. The intensity of WT inoculated with CMV-Δ2b was set as 1. Ponceau, 25S rRNA and U6 RNA were used as the loading controls. Blot-experiments per mutant line were repeated three times with similar results showed in Figure S6 (**B**-**G**). Relative viral protein accumulation (**E**), viral RNAs accumulation (**F**) and vsiRNAs accumulation (**G**) in WT and *ago2b* plants infected with CMV-Fny or CMV-Δ2b at 19 dpi. Values represent the average of nine individual blot-experiments results, three lines of *ago2b* with three technical replicates each. Error bars represent SD. ‘‘%’’ indicates the percentage of increased accumulation in *ago2b* compared to WT (100%).

We also further examined virus accumulation in these *ago2b* mutants compared to Wildtype Micro-Tom. Unexpectedly, Western blot or Northern blot results respectively showed that, after the infection of CMV-Δ2b or CMV-Fny, the accumulation of CP protein, viral RNAs or vsiRNAs was not affected in all three different *ago2b* mutants compared to wildtype Micro-Tom (Fig 3B-D and S6B-G). To further demonstrate accumulation variation of CP, viral RNAs and vsiRNAs in *ago2b* mutant, we also statistically calculated their relative accumulation based on three replicate results for each different *ago2b* mutant. It was found that, after the infection of CMV-Fny or CMV-Δ2b, viral CP, RNAs or vsiRNAs was not significantly changed in *ago2b* mutant compared to wildtype Micro-Tom (Fig 3E-G). These results indicate that, unlike AGO2a, AGO2b does not play major roles in antiviral RNAi in tomato.

### 4. AGO2a and AGO2b don’t function redundantly in antiviral defense and plant development in tomato

We further found that tomato *ago2ab* double mutants did not exhibit any defects in growth and development compared to *ago2a, ago2b* single mutants or wildtype Micro-Tom plants (Fig S4 and Fig 4A), indicating that AGO2a and AGO2b do not play redundant roles in regulating plant developments. To further find out whether AGO2a and AGO2b play redundant roles in antiviral defense, we infected *ago2ab-367, ago2ab-601* or *ago2ab-615* double mutants along with wildtype Micro-Tom plants with wildtype CMV-Fny or CMV-Δ2b, respectively. It was also found that none of three allelic *ago2ab* mutants show defects in growth and development after CMV-Δ2b infection (Fig 4A). And after wildtype CMV-Fny infection, these *ago2ab* mutants displayed disease symptoms similar to *ago2a, ago2b* single mutants or wildtype Micro-Tom plants (Fig 4A and S7A). Thus, AGO2a and AGO2b do not function redundantly to regulate disease symptom induction in tomato after viral infection either.

**Figure 4.**
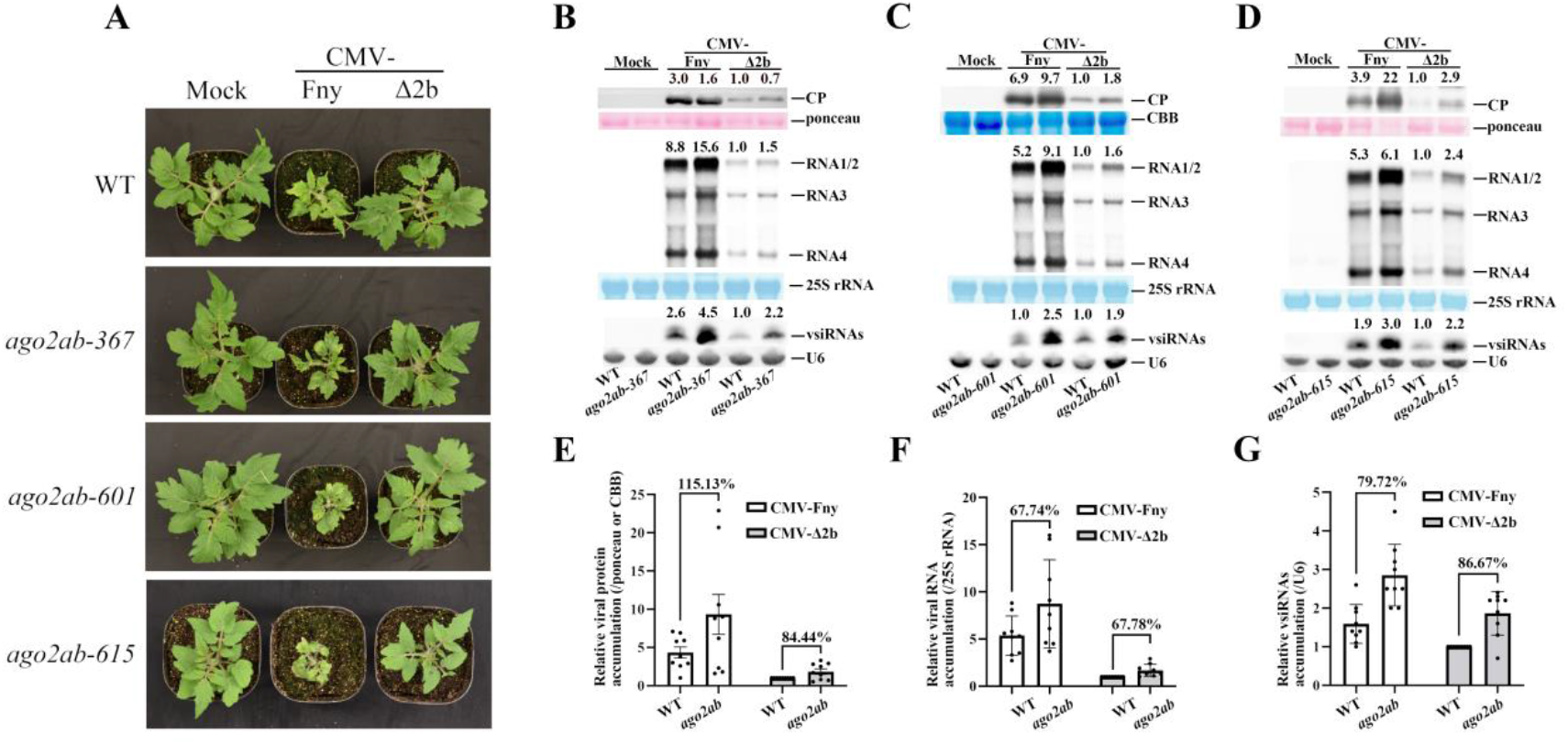
AGO2a and AGO2b don’t function redundantly in antiviral defense and plant development in tomato. (**A**) Wild-type (WT) and three lines of *ago2ab* mutants (*367, 601* and *615*) were photographed 19 days post-inoculation (dpi) with buffer (Mock), CMV-Fny or CMV-Δ2b. Symptoms induced with CMV-Fny were shown in Figure S7**A**. (**B**-**D**) Viral protein, viral RNAs and vsiRNAs accumulation in WT and three lines of *ago2ab* plants, infected with CMV-Fny or CMV-Δ2b at 19 dpi. Values at the top panel represent corresponding hybridization signal intensity of viral CP, RNAs 1-4, and vsiRNAs. The intensity of WT inoculated with CMV-Δ2b was set as 1. Ponceau, CBB, 25S rRNA and U6 RNA were used as the loading controls. Blot-experiments per mutant line were repeated three times with similar results showed in Figure S7 (**B**-**G**). Relative viral protein accumulation (**E**), viral RNAs accumulation (**F**) and vsiRNAs accumulation (**G**) in WT and *ago2ab* plants infected with CMV-Fny or CMV-Δ2b at 19 dpi. Values represent the average of nine individual blot-experiments results, three lines of *ago2ab* with three technical replicates each. Error bars represent SD. ‘‘%’’ indicates the percentage of increased accumulation in *ago2ab* compared to WT (100%).

We further examined CP protein or viral RNAs accumulation in *ago2ab* double mutants using Western blot or Northern blot. It was found that CP protein and viral RNAs were significantly increased in these *ago2ab* knockout mutants after the infection of either wildtype CMV-Fny or CMV-Δ2b, compared to that in wildtype Micro-Tom plants (Fig 4B-D up and S7B-G up). vsiRNAs accumulation in these *ago2ab* double mutants was also increased compared to wildtype Micro-Tom plants (Fig 4B-4D down and S7B-G down). To further demonstrate accumulation variation, we also statistically calculated relative accumulations of CP, viral RNAs and vsiRNAs in *ago2ab* mutant based on three replicate results for each different *ago2ab* mutant. It was found that, after the infection of CMV-Fny or CMV-Δ2b, viral CP was significantly increased by ∼115% or ∼84% in *ago2ab* mutant compared to wildtype Micro-Tom (Fig 4E), viral RNAs was significantly increased by ∼68% or ∼68% (Fig 4F), and vsiRNAs was significantly increased by ∼79% or ∼86% (Fig 4G). However, the relative accumulation of CP, viral RNAs or vsiRNAs in *ago2ab* double mutant was comparable to that in *ago2a* single knockout mutant (Fig 3E-G). These results indicate that only AGO2a plays important roles in antiviral RNAi, and AGO2a and AGO2b do not function redundantly in antiviral defense in tomato.

### 5. Expression pattern underlying the antiviral function of AGO2a and AGO2b in tomato

The distinct difference in antiviral defense between AGO2a and AGO2b prompts us to find the underlying mechanism. We then examined their transcription level for the two tandem-repeated homologs in tomato before and after viral infection. RT-PCR analysis indicated that *AGO2a* was substantially expressed in tomato plants and its expression was not affected in our *ago2a, ago2b* or *ago2ab* mutants compared to wildtype Micro-Tom plants (Fig 5A and S8A-C). Interestingly, we found that *AGO2a* mRNA expression was drastically elevated after wildtype CMV-Fny infection, though it was not clear after CMV-Δ2b infection (Fig 5A and S8A-B). However, *AGO2b* expression cannot be detected in these plants before or after infection of either wildtype CMV-Fny or CMV-Δ2b by RT-PCR analysis (Fig 5A and S8A-C). Further quantitative RT-PCR analysis demonstrated the same expression pattern of *AGO2a* and *AGO2b* in these plants, except that *AGO2a* and *AGO2b* mRNA expression was modestly increased in *ago2ab* mutants after CMV-Fny infection (Fig 5C-D).

**Figure 5.**
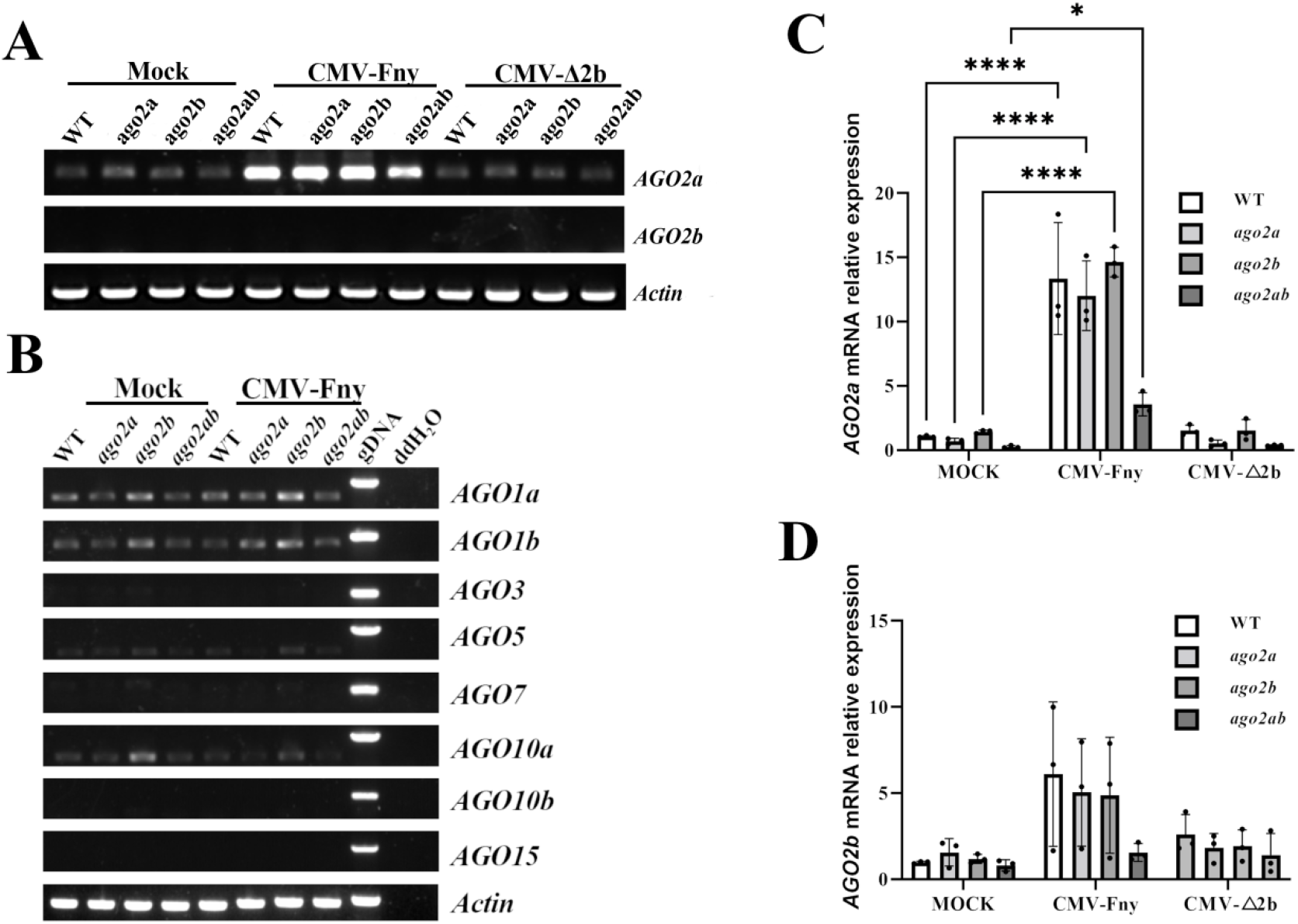
Expression of AGO genes in different genotypes after the infection of CMV-Fny or CMV-Δ2b. (**A**) Semi-quantitative RT-PCR analysis of *AGO2a* and *AGO2b* genes in WT, *ago2a, ago2b* and *ago2ab* in Micro-Tom. Experiments were repeated three times with similar results showed in Figure S6 (**A** and **B**). (**B**) Semi-quantitative RT-PCR analysis of AGO genes in WT, *ago2a, ago2b* and *ago2ab* with buffer (Mock) or CMV-Fny infection. Experiments were also repeated three times with similar results showed in Figure S8 (**D** and **E**). Relative *AGO2a* (**C**) or *AGO2b* (**D**) expression in mutant tomato plants inoculated with CMV-Fny or CMV-Δ2b at 19 dpi. Error bars represent SD. Experiments were repeated three times with similar results. Asterisks indicate a statistically significant difference (\**p*<0.01, \*\*\*\**p*<0.0001).

We also examined transcriptional expressions of other tomato AGOs. RT-PCR results showed that none of them was induced by the infection of CMV-Fny or CMV-Δ2b, and their expressions were not affected in *ago2a, ago2b* or *ago2ab* mutants (Fig 5B, S8D-E), indicating that *AGO2a* was the sole effector of antiviral RNAi which would be significantly induced to against viral infection in tomato plants.

We further cloned genomic fragments of *AGO2a* or *AGO2b* gene into expression vector pCambia-3301 to be fused with GFP in their C-terminals and driven by *ACTIN2* promoter (Fig 6A), then examined the subcellular localization of both GFP-fused AGO2a and AGO2b proteins. In transient expression, it was found that strong AGO2a-GFP signal were detected to co-localize with endo-membrane marker PIP2A in the leaves of *N. benthamiana* when observed under confocal microscope, while GFP signal expressed from empty vector are distributed in different subcellular structures including ER, cytosol and nuclei (Fig 6B). However, AGO2b-GFP signal was barely detected on endo-membrane compared to AGO2a-GFP (Fig 6B). We further found that AGO2a-GFP protein with correct 130KD molecular weight could be readily detected by Western blot, but AGO2b-GFP protein could not be detected in the transient expression (Fig 6C). Thus, these variations of expression pattern between AGO2a and AGO2b are well consistent with their distinct functions in antiviral RNAi in tomato, which probably underlies their roles in antiviral defense in tomato.

**Figure 6.**
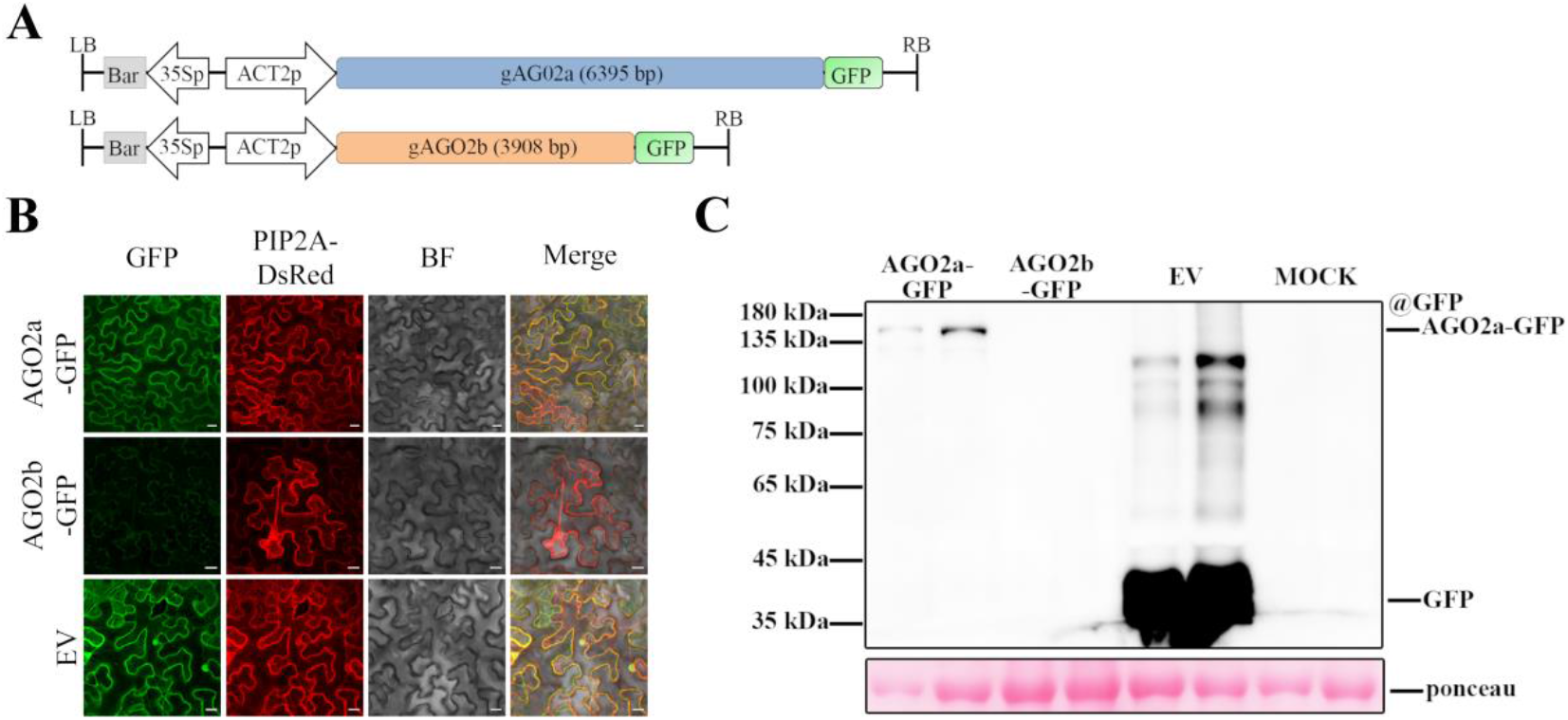
Variation of expression between AGO2a and AGO2b. (**A**) Schematic diagram to illustrate the vector pCambia-3301-gAGO2a/gAGO2b, used for AGO2a and AGO2b transient expression in the leaves of *N. benthamiana*. (**B**) Subcellular localization of AGO2a and AGO2b fused to GFP transiently expressed in *N. benthamiana*. Scale bar, 20μm. (**C**) Western blot analysis of AGO2a and AGO2b fused to GFP protein transiently expressed in *N. benthamiana* leaves.

## DISCUSSION

AGO2 is one key effector mediating antiviral RNAi immunity in plants (Harvey et al., 2011; Jaubert et al., 2011; Wang et al., 2011; Alazem et al., 2017). Compared to the single AGO2 in Arabidopsis (Tolia and Joshua-Tor, 2007), two differentiated AGO2 homologs, AGO2a and AGO2b, are evolved in tomato (Bai et al., 2012). In the research, we successfully generated *ago2a, ago2b* and *ago2ab* knockout mutants using CRISPR-Cas9 and infected them with wildtype CMV-Fny or VSR-deficient CMV, CMV-Δ2b, to investigate their antiviral function in tomato. It was found that only AGO2a was induced to function in the antiviral defense in tomato, AGO2b should be a pseudogene without function due to the diminished expression level of AGO2b in tomato. Thus, for the first time our research revealed the distinct function of two differentiated AGO2 effectors in antiviral immunity in tomato.

Interestingly, we found that AGO2a could defend against the infection of not only VSR-deficient CMV but also wildtype CMV-Fny in tomato, which is different from previous findings in Arabidopsis in which wildtype CMV-Fny can be efficiently replicated and propagated in both wildtype Arabidopsis plants and antiviral RNAi-defective mutants because VSR 2b almost completely inhibits antiviral RNAi in wildtype Arabidopsis plants. Therefore, our findings here indicated that antiviral RNAi immunity evolves to against the infection of wildtype virus with potent VSR in tomato. This function of antiviral RNAi may be shared in varieties of other plants.

In addition, our results showed that only wildtype CMV-Fny but not VSR 2b-deficient CMV (CMV-Δ2b) caused disease symptom in tomato plants, indicating the critical roles of 2b in disease symptom induction. Since wildtype CMV-Fny was abundantly accumulated much higher in tomato plants than CMV-Δ2b, it could be that viral accumulation exceeded threshold so that plants cannot maintain regular homeostasis for normal plant growth and development. However, it cannot be excluded that 2b may cause plant disease symptoms by directly interfering growth and developmental processes in tomato.

Surprisingly, we found that just like wildtype Micro-Tom plants, *ago2a, ago2b* or *ago2ab* mutants did not show disease symptoms after CMV-Δ2b infection, although virus accumulation was significantly increased in these mutants compared to wildtype Micro-Tom plants. And after wildtype CMV-Fny infection, *ago2a, ago2b* or *ago2ab* mutants did not show enhanced disease symptoms compared to wildtype Micro-Tom plants either, although viral accumulation was also dramatically increased in these mutants. These results suggested that, unlike in Arabidopsis, AGO2-mediated antiviral RNAi mainly inhibits viral accumulation but does not play a major role in preventing disease induction in tomato.

Therefore, our findings together show that novel mechanisms and specialized function of antiviral RNAi may be developed to counter viral aggression in tomato. It is probable that during the arm race between host and viruses, tomato plants gain new arsenals to attenuate the inhibition of VSR on antiviral RNAi, so that antiviral RNAi in tomato maintains to counteract the infection and propagation of wildtype viruses. However, novel mechanisms in tomato rather than AGO2-mediated antiviral RNAi may play major roles in promoting plant tolerance to maintain health after viral infection. It will be very interesting to find these detailed mechanisms in the future.

## MATERIALS AND METHODS

### Viruses and Plant materials

Wild-type virus CMV-Fny is a strain of CMV subgroup 1, isolated and cloned from a muskmelon farm in New York (Rizzo and Palukaitis, 1990). CMV-Δ2b is a mutant virus of CMV-Fny. In CMV-Δ2b, the third codon UUG in 2b ORF encoded by the CMV-Fny is mutated to stop codon UAG, and three AUG codons at the first, 8th and 18th positions in 2b ORF were mutated to ACG so that the amino acids encoded by the overlapping part of the 2a ORF were not changed (Wang et al., 2011).

Tomato cv. Micro-Tom (MT) was used as wild types in this study. *ago2a, ago2b* and *ago2ab* knockout mutants were generated in MT background by CRISPR/Cas9 genome editing. All seedlings used for virus inoculations were grown in an insect-free growth room under 24°C with a 12 h light/12 h in dark photoperiod, light intensity of 8000 lux.

### Construction of genome editing vector

Two sequences (Solyc02g069260 and Solyc02g069270) of tomato *AGO2* gene were identified using the Sol genomics network database (Fernandez-Pozo et al., 2015) according to the reported tomato AGO2 sequence (Bai et al., 2012). Target sites of these two sequences for editing tomato AGO2 genome were selected using the online tool CCTop-CRISPR/Cas9 target online predictor and Cas-OFFinder (Bae et al., 2014; Stemmer et al., 2015). The pHEE401 vectors (Fig. S2) are used to generate homozygous mutants for two target genes in Arabidopsis with high efficiency (Wang et al., 2015). To construct pHEE401-AGO2a-sgRNA, pHEE401-AGO2b-sgRNA and pHEE401-AGO2ab-sgRNA binary vector, target-specific sgRNA expression cassettes were cloned into the pHEE401 backbone. Briefly, fragments of the sgRNA expression cassettes were amplified from pCBC-DT1T2_tomatoU6 with primer pairs listed in supplementary Table S1, and inserted into pHEE401 vectors using BsaI restriction enzyme site. Subsequently, these three recombinant vectors were each transformed by heat shock into the *A. tumefaciens* strain GV3101. Primers used in this study were listed in supplementary Table S1.

### Tomato transformation

Agrobacterium-mediated transformations of tomato cotyledons were performed to generate *ago2a, ago2b* and *ago2ab* knockout transgenic tomato plants, as described in previous research (Sun et al., 2006). Briefly, cotyledon segments from aseptically seedlings were placed on Murashige and Skoog (MS) medium, and precultured in the dark for 2-day. Then, Cotyledon explants were soaked in MS liquid medium containing Agrobacterium for 10 min, and also co-cultivated on MS medium in dark for 2-day. Following, these cotyledon explants were transferred to a callus induction medium containing 75 mg L^−1^ kanamycin to select transgenic cells. When small shoot buds were induced from callus, they were transferred to shoot elongation medium containing 50 mg L^−1^ kanamycin. Shoots (approximately 1.5 cm tall) then were excised from shoot buds and inserted to rooting medium without hormones for root regeneration. Finally, well-rooted plants were planted in greenhouse under 26°C with a 16 h light/8 h in dark photoperiod, light intensity of 20000 lux.

### DNA Extraction and Mutant identification

To detect gene editing, genomic DNA was extracted from tomato leaves using a CTAB method as described in previous research (Doyle, 1987). The DNA fragments containing target sites were amplified by PCR with primer pairs listed in supplementary Table S1. The PCR products were then sequenced by Sanger sequencing to analysis mutation. Positive mutant plants (without Cas9) were planted in greenhouse under 26°C with a 16 h light/8 h in dark photoperiod, light intensity of 20000 lux.

### RNA Extraction and Northern Blot Analysis

To analyze viral RNA accumulation, systemically infected leaves were collected at 19 days post-inoculation, and each sample collected from three plants. RNA extraction and Northern blots were conducted according to the published protocol (Guo et al., 2019b). Ten μg of total or small RNAs was loaded each lane for Northern analysis of the viral genomic or vsiRNAs accumulation. Northern blots were performed with probes of Biotin-dUTP labelled cDNA or DNA oligonucleotides, as described previous research (Wang et al., 2011). The blot signal was detected by chemiluminescence image analysis system (Tanon-5200).

### Protein Extraction and Western Blot Analysis

Total proteins were extracted from leaf samples according to method described in published research (Wang et al., 2003). Equal amounts of proteins were transferred to PVDF membranes after being separated on 10% or 12.5% SDS-PAGE gels. Viral protein was detected using rabbit polyclonal anti-CP antibody (1:3000; Zoonbio Biotechology) specific to CMV-Fny CP. GFP -fused proteins were detected using rabbit monoclonal anti-GFP antibody (1:3000; Abcam). The blot signal was detected by chemiluminescence image analysis system (Tanon-5200).

### RT-PCR and quantitative real-time RT-PCR (qRT-PCR) analysis

cDNA was synthesized using the HiScript II cDNA Synthesis Kit (Vazyme) according to manufacturer’s instructions. AGO genes were subjected to semi-quantitative RT-PCR, and products of 28 cycles were segregated in 1.0% agarose gel. qRT-PCR was performed using Taq Pro Universal SYBR qPCR Master Mix (Vazyme). Tomato actin gene was used as an internal control and normalizers. Primers used in this study were listed in supplementary Table S1. All experiments were repeated three times.

### Observation of protein subcellular localization

For subcellular localization, tomato *AGO2a* and *AGO2b* genomic DNA were amplified with primer pairs listed in supplementary Table S1. DNA sequences were integrated into p3301vectors between the SpeI and SmaI sites and each transformed by heat shock into the *A. tumefaciens* strain GV3101. *A. tumefaciens* cells, harboring p3301-EGFP, p3301-EGFP-AGO2a or p3301-EGFP-AGO2b, were infiltrated separately with PIP2A-DsRed (membrane marker) into the fifth or sixth true leaves of *N. benthamiana*. The final densities of *A. tumefaciens* cells were equivalent to an A600 of 0.5. Leaves were examined for GFP signal at 36-h post agroinfiltration by fluorescence microscopy (Leica DMI 6000B with filter block L5 containing a 480/40 nm excitation filter, a 505 nm dichroic mirror and a 527/30 nm barrier filter for GFP fluorescence), and images were taken using Leica LAS AF software.

### Accession numbers

Sequence data used in this study are downloaded from Sol Genomics ([Sol Genomics Network](https://solgenomics.net/)). The accession numbers are as follows: *AGO2a* (Solyc02g069260), *AGO2b* (Solyc02g069270), *AGO1a* (Solyc06g072300), *AGO1b* (Solyc03g098280), *AGO3* (Solyc02g069280), *AGO5* (Solyc06g074730), *AGO*6 (Solyc07g049500), *AGO10*a (Solyc09g082830), *AGO10b* (Solyc12g006790), *AGO15* (Solyc03g111760), *Actin* (Solyc11g005330), *NtAGO2* (XM_016629769), *AtAGO2* (AT1G31280), *OsAGO*2 (XP_015636011).

## ACKNOWLEDGEMENTS

The authors thank professor Chuanyou Li sharing the construct pHEE401, and Chuanlong Sun, Yuzhen Mei for experimental advice; and all members in Zhongxin Guo’s and Yanhong Han’s groups for helpful suggestions. This work was supported by National Natural Science Foundation of China (31870146 and 32160619), the Science Foundation of Fujian province (2020J02014) and “Hundred Talent” of Fujian Province.

## AUTHOR CONTRIBUTIONS

Z.X.G. and L.L.Z. designed the experiments, analyzed the data, and wrote the manuscript. L.L.Z., Y.F.C., X.M.X, H.Y.G., and J.M.C. performed the experiments. All authors read and approved of the manuscript.

