## Supplementary material for "AGO2a but not AGO2b mediates antiviral defense against the infection of wildtype Cucumber mosaic virus in tomato": Sup Figures

**Figure S1. Protein structures and similarity of amino acid sequences of AGO2 proteins.**

(**A**) Schematic diagrams were illustrated protein structures of the AGO2, originated from tomato, tobacco, Arabidopsis and rice. Protein structures of AGO2 were produced using the online tool SMART (http://smart.embl.de). (**B**) Similarity of amino acid sequences of AGO2a with other AGO2s. Multiple sequence alignments were performed using MegAlign suite included in DNASTAR software version 11.

**Figure S2. Multiple alignment of AGO2 proteins originated from tomato, tobacco, Arabidopsis and rice.**

The major functional domains of AGO2 are indicated by colored boxes.

**Figure S3. Multiple alignment of ArgoN domain of AGO2a proteins.**

Rectangular box indicates the deleted region only in AGO2b, but retained in AGO2a, NtAGO2, AtAGO2 and OsAGO2.

**Figure S4. ago2a, ago2b and ago2ab do not show defects in growth and development compared to WT Micro-tom.**

**Figure S5. *ago2a mutants are defective in antiviral* immunity.**

(**A**) Wild-type (WT) and three lines of *ago2a* mutants (*126*, *338* and *453*) were photographed 19 days post-inoculation (dpi) with CMV-Fny. Accumulation of viral protein, RNAs and vsiRNAs in WT and mutant lines of *ago2a-126* (**B**, **E**), *ago2a-338* (**C**, **F**) and *ago2a-453* (**D**, **G**). Blot-experiments per mutant line were repeated three times. The figure above shows the results of two additional experimental replicates. Pnoceau, 25S rRNA and U6 RNA were used as the loading controls. Values at the top panel represent corresponding hybridization signal intensity of viral CP, RNAs 1-4, and vsiRNAs. The intensity of WT inoculated with CMV-∆2b was set as 1.

**Figure S6. *AGO2b* does not function in antiviral immunity in tomato.**

(**A**) Wild-type (WT) and three lines of *ago2b* mutants (*281*, *293* and *620*) were photographed 19 dpi with CMV-Fny. Accumulation of viral protein, RNAs and vsiRNAs in WT and mutant lines of *ago2b-281* (**B**, **E**), *ago2b-293* (**C**, **F**) and *ago2b-620* (**D**, **G**). Blot-experiments per mutant line were repeated three times. The figure above shows the results of two additional experimental replicates. Ponceau, 25S rRNA and U6 RNA were used as the loading controls. Values at the top panel represent corresponding hybridization signal intensity of viral CP, RNAs 1-4, and vsiRNAs. The intensity of WT inoculated with CMV-∆2b was set as 1.

**Figure S7. AGO2a and AGO2b do not function redundantly in antiviral defense in tomato.**

(**A**) Wild-type (WT) and three lines of *ago2ab* mutants (*367*, *601* and *615*) were photographed 19 days post-inoculation (dpi) with CMV-Fny. Accumulation of viral protein, RNAs and vsiRNAs in WT and mutant lines of *ago2ab-367* (**B**, **E**), *ago2b-601* (**C**, **F**) and *ago2ab-615*(**D**, **G**). Blot-experiments per mutant line were repeated three times. The figure above shows the results of two additional experimental replicates. Ponceau, CBB, 25S rRNA and U6 RNA were used as the loading controls. Values at the top panel represent corresponding hybridization signal intensity of viral CP, RNAs 1-4, and vsiRNAs. The intensity of WT inoculated with CMV-∆2b was set as 1.

**Figure S8. Expression of AGO genes in different genotypes inoculated with CMV-Fny or CMV-Δ2b.**

(**A** and **B**) Semi-quantitative RT-PCR analysis of *AGO2a* and *AGO2b* genes in WT, *ago2a*, *ago2b* and *ago2ab* in Micro-Tom. The figure above shows the results of two additional experimental replicates. (**C**) PCR detection of *AGO2b* and *AGO2b* genes in Micro-Tom cDNA and genomic DNA (gDNA). PCR products of *AGO2b* were obtained from gDNA, suggesting that lack of PCR products from cDNA is due to lower gene expression instead of failure of PCR. (**D** and **E**) Semi-quantitative RT-PCR analysis of AGO genes in WT, *ago2a*, *ago2b* and *ago2ab* with buffer (Mock) or CMV-Fny infection. The figure above shows the results of two additional experimental replicates.

Table S1. Primers used in this study

**Function**

**Primer name**

**Sequence 5' to 3'**

**Reference**

construction of pHEE401 binary vector, used to knockout *AGO2a* gene

*AGO2a*-DT1-BsF

ATATATGGTCTCGATTGATCGTGGGAACTACCGACGGTT

In this study

*AGO2a*-DT1-F0

TGATCGTGGGAACTACCGACGGTTTTAGAGCTAGAAATAGC

In this study

*AGO2a*-DT2-R0

AACCCACGGCCTTGTCCTCGACCAATCTCTTAGTCGACTCTAC

In this study

*AGO2a*-DT2-BsR

ATTATTGGTCTCGAAACCCACGGCCTTGTCCTCGACCAA

In this study

construction of pHEE401 binary vector, used to knockout *AGO2b* gene

*AGO2b*-DT1-BsF

ATATATGGTCTCGATTGGTGGAGCTCCGCCGGGTCGGTT

In this study

*AGO2b*-DT1-F0

TGGTGGAGCTCCGCCGGGTCGGTTTTAGAGCTAGAAATAGC

In this study

*AGO2b*-DT2-R0

AACTTGCCAAGCGTTACCACCACAATCTCTTAGTCGACTCTAC

In this study

*AGO2b*-DT2-BsR

ATTATTGGTCTCGAAACTTGCCAAGCGTTACCACCACAA

In this study

construction of pHEE401 binary vector, used to knockout *AGO2a* and *AGO2b* gene

*AGO2a-*DT1-BsF

ATATATGGTCTCGATTGGTCTGACGGGGAAGATGCTGTT

In this study

*AGO2a*-DT1-F0

TGGTCTGACGGGGAAGATGCTGTTTTAGAGCTAGAAATAGC

In this study

*AGO2b*-DT2-R0

AACCGTTGGTAGTTTCCACGGTCAATCTCTTAGTCGACTCTAC

In this study

*AGO2b*-DT2-BsR

ATTATTGGTCTCGAAACCGTTGGTAGTTTCCACGGTCAA

In this study

Genotyping for *ago2a* mutants

*AG02a*-detect-F

CGAGTTGTTTACTCCTAAG

In this study

*AG02a*-detect-R

CGTTGAACCGGTTGGTTTAC

In this study

Genotyping for *ago2b* mutants

*AG02b*-detect-F

GACCGTGGAAACTACCAACG

In this study

*AG02b*-detect-R

CCAGAAGATTGAGGACGATC

In this study

Genotyping for *ago2ab* mutants

*AG02ab*-2a-detect-F

GATCAGAGTCCATCATCTCG

In this study

*AG02ab*-2a-detect-R

GACCTACAGAAGTCCTGCAC

In this study

*AG02ab*-2b-detect-F

GAGTCAGAAGTGTATACAAC

In this study

*AG02ab*-2b-detect-R

CAAGCAGTGATTGGAACACC

In this study

using for semi-quantitative RT-PCR analysis of *AGO2a*

*AGO2a*-F

TTGGCGAAGGCTATATACGACAG

Miao et al., 2012

*AGO2a*-R

AGTTGCAGAGCTAGGAGAGTTCATC

Miao et al., 2012

using for semi-quantitative RT-PCR analysis of *AGO2b*

*AGO2b*-F

ATCGTTACAAGTATAAACCTGAAATCAC

Miao et al., 2012

*AGO2b*-R

GGCAGGTGAAGTTGTAGAGCTAGAA

Miao et al., 2012

using for semi-quantitative RT-PCR analysis of *AGO1a*

*AGO1a*-F

CTATCAGCCCCCAGTTACGTTTG

Miao et al., 2012

*AGO1a*-R

ATCACCCTTTTGACATTCTCCTTG

Miao et al., 2012

using for semi-quantitative RT-PCR analysis of *AGO1b*

*AGO1b*-F

CAAGACACTGTTGCACATGGGTT

Miao et al., 2012

*AGO1b*-R

CAAAAATGATTGTAGGACGGTCG

Miao et al., 2012

using for semi-quantitative RT-PCR analysis of *AGO3*

*AGO3*-F

CACAGAAATCCAGGACTGAAAAGA

Miao et al., 2012

*AGO3*-R

CGTTGTAAATCGACAGATATGAAGC

Miao et al., 2012

using for semi-quantitative RT-PCR analysis of *AGO5*

*AGO5*-F

TAGTAAGAACATGCCTTTCCTCACC

Miao et al., 2012

*AGO5*-R

GCAGCTCTTACATCCACGTCATTC

Miao et al., 2012

using for semi-quantitative RT-PCR analysis of *AGO7*

*AGO7*-F

GGCTGGCATGTTCGAGATTTC

Miao et al., 2012

*AGO7*-R

GTGGCACCAGTGAAATAGGCTTG

Miao et al., 2012

using for semi-quantitative RT-PCR analysis of *AGO10a*

*AGO10a*-F

CGAGTTAGATGCAATTAGGAAGGC

Miao et al., 2012

*AGO10a*-R

AATTAGTTTCAGGCATGTCTGGTTC

Miao et al., 2012

using for semi-quantitative RT-PCR analysis of *AGO10b*

*AGO10b*-F

CCAGGAGGTGTATGGATACACC

Miao et al., 2012

*AGO10b*-R

CTCGAGCTTCCACTGATGCAAGC

Miao et al., 2012

using for semi-quantitative RT-PCR analysis of *AGO15*

*AGO15*-F

TCTCTTTAGACCAGTTTCAGATAGGG

Miao et al., 2012

*AGO15*-R

CATCTGAGCTGAAACCAAACGAG

Miao et al., 2012

using for qRT-PCR analysis of *AGO2a*

qPCR-*AGO2a*-F

TACCTTCAGGAACAACATTCGA

Miao et al., 2012

qPCR-*AGO2a*-R

GGGATCTTCAAGGGGAAAAGTA

Miao et al., 2012

using for qRT-PCR analysis of *AGO2b*

qPCR-*AGO2b*-F

GTCACGCACTTATGACATTACC

Miao et al., 2012

qPCR-*AGO2b*-R

CTTGCAGTATATCACGAGGAGT

Miao et al., 2012

using for semi-quantitative RT-PCR and qRT-PCR analysis of *ACTIN*

qPCR-actin-F

GGTCCTCTTCCAGCCATCC

Miao et al., 2012

qPCR-actin-R

CCACTGAGCACAATGTTACCG

Miao et al., 2012

construction of p3301 binary vector, used to observe *AGO2a* subcellular localization

p3301-*AGO2a*-GFP-F

ATCGTGTGTGACCTCGAGACTAGTATGGATCGTGGGAACTACCGAC

In this study

p3301-*AGO2a*-GFP-R

ACCAGGTGGAGGTCCCCCGGGGACGAAAAACATTACGTTCTGC

In this study

construction of p3301 binary vector, used to observe *AGO2b* subcellular localization

p3301-*AGO2b*-GFP-F

ATCGTGTGTGACCTCGAGACTAGTATGGACCGTGGAAACTACCAAC

In this study

p3301-*AGO2b*-GFP-R

ACCAGGTGGAGGTCCCCCGGGGACAAAAAACATAATGTTCTGC

In this study
